# Mutation load decreases with haplotype age in wild Soay sheep

**DOI:** 10.1101/2021.03.04.433700

**Authors:** M.A. Stoffel, S.E. Johnston, J.G. Pilkington, J.M Pemberton

## Abstract

Runs of homozygosity (ROH) are pervasive in diploid genomes and expose the effects of deleterious recessive mutations, but how exactly these regions contribute to variation in fitness remains unclear. Here, we combined empirical analyses and simulations to explore the deleterious effects of ROH with varying genetic map lengths in wild Soay sheep. Using a long-term dataset of 4,592 individuals genotyped at 417K SNPs, we found that inbreeding depression increases with ROH length. A 1% genomic increase in long ROH (>12.5cM) reduced the odds of first-year survival by 12%, compared to only 7% for medium ROH (1.56-12.5cM), while short ROH (<1.56cM) had no effect on survival. We show by forward genetic simulations that this is predicted: compared with shorter ROH, long ROH will have higher densities of deleterious alleles, with larger average effects on fitness and lower population frequencies. Taken together, our results are consistent with the idea that the mutation load decreases in older haplotypes underlying shorter ROH, where purifying selection has had more time to purge deleterious mutations. Finally, our study demonstrates that strong inbreeding depression can persist despite ongoing purging in a historically small population.

## Introduction

The structure of deleterious genetic variation in natural populations shapes a range of processes in evolutionary biology, such as the strength of inbreeding depression and the efficiency of genetic purging (Charlesworth & Willis, 2009; Hedrick & Garcia-Dorado, 2016). The role of deleterious mutations is also increasingly discussed in applied conservation, in particular when considering genetic rescue of small populations (Kyriazis, Wayne, & Lohmueller, 2020; Ralls, Sunnucks, Lacy, & Frankham, 2020). To date, studies in wild populations have mostly focused on average, genomewide fitness effects of deleterious recessive alleles through measuring genome-wide inbreeding coefficients (Bérénos, Ellis, Pilkington, & Pemberton, 2016; Chen, Cosgrove, Bowman, Fitzpatrick, & Clark, 2016; Harrisson et al., 2019; Hoffman et al., 2014; Huisman, Kruuk, Ellis, Clutton-Brock, & Pemberton, 2016; Niskanen et al., 2020), or genome-sequence based predictions of deleterious mutations (Grossen, Guillaume, Keller, & Croll, 2020; Robinson, Brown, Kim, Lohmueller, & Wayne, 2018; Xue et al., 2015). Therefore, we still know very little about how deleterious mutations in different parts of the genome contribute to inbreeding depression, as these analyses usually require large samples of individuals with known fitness and dense genomic data – both of which are scarce in wild non-model organisms.

In populations for which mapped genetic markers are available, runs of homozygosity (ROH) open up new possibilities for studying the effects of (partially) recessive deleterious variation. These long stretches of homozygous genotypes are ubiquitous in diploid genomes and commonly arise when individuals inherit homologous haplotypes which are identical-by-descent (IBD), originating from a single copy of the region in a common ancestor. Offspring of related parents have more ROH which in turn increases the probability that partially recessive deleterious alleles are expressed, thereby causing inbreeding depression (Charlesworth & Willis, 2009). The lengths and numbers of ROH can vary widely between individuals, and have been shown to contribute to the genetic architecture of complex traits and diseases in humans (Ceballos, Joshi, Clark, Ramsay, & Wilson, 2018; Clark et al., 2019) and to production traits in livestock (Ferenčaković, Sölkner, Kapš, & Curik, 2017; Pryce, Haile-Mariam, Goddard, & Hayes, 2014). In wild populations, ROH are increasingly used to precisely measure individual inbreeding coefficients (Kardos et al., 2018; Kardos, Luikart, & Allendorf, 2015) and the effects of inbreeding on fitness (Bérénos et al., 2016; Stoffel, Johnston, Pilkington, & Pemberton, 2020). Moreover, genome-wide association studies are starting to uncover associations between ROH at specific locations in the genome and complex traits or fitness, thereby providing information about the distribution of effect sizes at loci causing inbreeding depression (Pryce et al., 2014; Stoffel et al., 2020).

The length of an ROH allows one to estimate the time to a most recent common ancestor (MRCA) of the underlying IBD haplotypes (Thompson, 2013). In any given generation, DNA is inherited in physically large chunks with genetic map lengths of around 100 cM, and recombination breaks up these segments in successive generations. For example, an initial segment is broken up into smaller IBD segments with an expected length of 2 cM after 25 generations, or 50 meioses (Thompson, 2013). The expected genetic map length *L* of an ROH can be estimated as *L* = 100/(2\**g*) cM, where *g* is the number of generations to the MRCA (Thompson, 2013), though the distribution is exponential with high variance due to stochastic effects of recombination and Mendelian segregation (Kardos, Taylor, Ellegren, Luikart, & Allendorf, 2016; Thompson, 2013). Long ROH originating from close inbreeding are expected to have a recent ancestor, while short ROH have an ancestor further back in the pedigree. In addition, when the effective size N_e_ of a population has been small at a given point in the recent history, ROH with a MRCA at that point will be more abundant in the current population. The relative frequencies of ROH of different lengths are therefore informative about recent fluctuations in population sizes (Browning & Browning, 2015; Ceballos et al., 2018; Kardos, Qvarnström, & Ellegren, 2017).

Considering jointly the fitness effects and sizes of ROH allows us to investigate how ROH lengths (and therefore haplotype ages) are associated with inbreeding depression and mutation load. Given that ROH lengths provide an expectation for the number of generations for which the underlying haplotypes have been exposed to selection, we hypothesise the following: Short ROH originating further back in the pedigree should be depleted of deleterious recessive variation, as purifying selection has had many generations to remove these mutations. In contrast, long ROH emerging from younger haplotypes should on average carry larger numbers of strongly deleterious recessive mutations at lower frequencies, and therefore be associated with stronger effects on fitness. In humans, ROH in general and especially long ROH are enriched for mutations which are predicted to be deleterious (Pemberton & Szpiech, 2018; Szpiech et al., 2018, 2013), but to our knowledge, these predictions have not been tested using actual fitness data in a wild population. Quantifying the fitness effects of different ROH length classes could help to understand the genetic basis of inbreeding depression and provide a novel way to assess the efficiency of selection against deleterious mutations in wild populations.

Here, we combined long-term life-history data for 4789 wild Soay sheep with 417K SNP genotypes and linkage map information to test whether inbreeding coefficients (F_ROH_) calculated from ROH with long, medium and short genetic map lengths differ in their contribution to inbreeding depression in first-year survival. We then used forward genetic simulations based on the Soay sheep demographic history to quantify the expected differences in the mutation load among ROH length classes and to explore the underlying causes. We discuss how our results fit into current knowledge about inbreeding depression and purging in small populations and methodological implications for studying the genetic basis of inbreeding depression.

## Materials and Methods

### Study population

Soay sheep are descendants of primitive European domestic sheep and have lived unmanaged on the St. Kilda archipelago, Scotland, for thousands of years (Clutton-Brock & Pemberton, 2004). A part of the population in the Village Bay area on the island of Hirta (57 49’N, 8 34’W) has been the focus of a long-term individual-based study since 1985 (Clutton-Brock & Pemberton, 2004). More than 95% of individuals in the study area are ear-tagged within a week after birth during the lambing season from March to May, and DNA samples are obtained from either blood samples or ear punches. Routine mortality checks, in particular during peak mortality at the end of winter, usually find around 80% of deceased animals (Bérénos et al., 2016). Here, we focused on the fitness trait ‘first year survival’, where every individual was given a 1 if it survived from birth (March to May) to the 30^th^ April of the next year, and a 0 if it did not, with measures available for 4879 individuals born from 1979 to 2018. In order to impute genotypes, we assembled a pedigree based on 438 unlinked SNP markers from the Ovine SNP50 BeadChip using the R package Sequoia (Huisman, 2017). In the few cases where no SNP genotypes were available, we used either observations from the field or microsatellite markers (Morrissey et al., 2012). All animal work was carried out according to UK Home Office procedures and was licensed under the UK Animals (Scientific Procedures) Act of 1986 (Project License no. PPL70/8818).

### Genotyping

We genotyped a total of 7,700 Soay sheep on the Illumina Ovine SNP50 BeadChip resulting in 39,368 polymorphic SNPs after filtering for SNPs with minor allele frequency > 0.001, SNP locus genotyping success > 0.99 and individual genotyping success > 0.95. We then used the check.marker function in GenABEL version 1.8-0 (Aulchenko, Ripke, Isaacs, & Van Duijn, 2007) with the same thresholds, including identity by state with another individual < 0.9. We also genotyped 189 sheep on the Ovine Infinium HD SNP BeadChip, resulting in 430,702 polymorphic SNPs for 188 individuals, after removing monomorphic SNPs, and filtering for SNPs with SNP locus genotyping success > 0.99 and individual sheep with genotyping success > 0.95. These sheep were specifically selected to maximise the genetic diversity represented in the full population (for full details, see Johnston, Bérénos, Slate, & Pemberton, 2016). All SNP positions were based on the Oar_v3.1 sheep genome assembly (GenBank assembly ID GCA_000298735.1 (Jiang et al., 2014)).

### Genotype imputation

The detailed genotype imputation methods are presented elsewhere (Stoffel et al., 2020). Briefly, we first merged the datasets from the 50K SNP chip and from the HD SNP chip with --bmerge in PLINK v1.90b6.12 (Purcell et al., 2007), resulting in a dataset with 436,117 SNPs including 33,068 SNPs genotyped on both SNP chips. We then discarded SNPs on the X chromosome and focused on the 419,281 SNPs located on autosomes. To impute SNPs with genotypes missing in individuals genotyped at the lower SNP density, we used AlphaImpute v1.98 (Hickey, Kinghorn, Tier, van der Werf, & Cleveland, 2012), which uses both genomic and pedigree information for phasing and subsequent imputation of missing genotypes. After imputation, we filtered SNPs with call rates below 95%. Overall, this resulted in a dataset with 7691 individuals, 417,373 SNPs and a mean genotyping rate per individual of 99.5% (range 94.8%-100%). We evaluated the accuracy of genotype imputation using 10-fold leave-one-out cross-validation. In each iteration, we randomly chose one individual genotyped on the high-density (HD) SNP chip, masked genotypes unique to the HD chip and imputed the masked genotypes. This allowed us to compare the imputed genotypes to the true genotypes and to evaluate the accuracy of the imputation. Overall, 99.3% of genotypes were imputed correctly. Moreover, the distribution of inbreeding coefficients F_ROH_ was very similar for individuals genotyped on the HD chip and individuals with imputed SNPs, indicating little difference in inferred ROH between the two groups and hence no obvious bias in ROH calling based on imputed genotypes (Stoffel et al., 2020).

### Inferring linkage map positions

We used a dense, sex-averaged Soay sheep linkage map with 36,972 autosomal markers (Johnston et al., 2016) to infer the genetic map positions in cM for each SNP in the imputed dataset. As the imputed SNP dataset used here had a higher SNP density than the linkage map SNP dataset, we interpolated the genetic positions of SNPs that were not present in the linkage map dataset by assuming a constant recombination rate in genomic regions between linkage mapped SNPs (Kardos et al., 2018, 2017). If two flanking SNPs had the same coordinates on the genetic map, all imputed SNPs in between were assigned the same genetic map position. If two flanking SNPs had different genetic map positions, the SNPs in between were assigned increasing genetic map positions depending on the physical distance to each of the two SNPs. For example, if the two flanking SNPs had cM positions 3 and 4, an imputed SNP half way between these SNPs on the physical map got assigned a cM position of 3.5. Imputed SNPs occurring before the first linkage mapped SNP on a chromosome were assigned a genetic map position of 0 cM, and SNPs occurring after the last linkage mapped SNP on a chromosome were assigned the same genetic position as the last linkage mapped SNP.

### ROH calling and individual inbreeding coefficients F_ROH_

We focused on ROH quantified by their genetic map lengths rather than physical map lengths as this accounts for the effects of recombination rate variation on detected ROH lengths (Kardos et al., 2017) and allows us to infer an expected time of coalescence for each ROH more precisely, assuming ROH are true IBD segments (Thompson, 2013). To call ROH based on their genetic map lengths in cM, we used PLINK (Purcell et al., 2007), replacing physical with genetic map positions in the input .*map* file. To keep the parameter arguments on a comparable scale to running PLINK with base-pair positions, we multiplied cM positions by 1e6. We then called ROH with a minimum length of 0.39 cM containing at least 25 SNPs while allowing a maximum gap of 0.25 cM between SNPs and one heterozygote genotype per ROH using the command ‘--homozyg --homozyg-window-snp 25 --homozyg-snp 25 --homozyg-kb 390 --homozyg-gap 250 --homozyg-density 100 --homozyg-window-missing 2 -- homozyg-het 2 --homozyg-window-het 2’. We semi-arbitrarily chose 0.39 cM as the minimum ROH length, which is the expected length of an ROH when the underlying haplotypes have a MRCA 128 generations ago as calculated with 100/(2g) cM (Thompson, 2013). Based on our SNP density, a stretch of genome with length 0.39 cM will contain on average ~50 SNPs, which, together with a slow LD decay in Soay sheep (Stoffel et al., 2020) should be sufficient to reliably call ROH of that length and above. To capture biologically interesting time-horizons, we qualitatively assessed the distribution of ROH lengths in the population (Supplementary Figure 1), and subsequently clustered ROH into three length classes: long ROH (> 12.5 cM) with an expected MRCA up to 4 generations ago and therefore likely to have originated from close inbreeding, medium ROH (1.56 – 12.5 cM) originating between 4 and 32 generations ago and reflecting the recent demographic history of the population and short ROH (0.39 – 1.56 cM) with an expected MRCA between 32 and 128 generations ago, reflecting deeper processes in the population history. For each length class, we calculated individual inbreeding coefficients F_ROH_ by summing up their total ROH length in each individual and dividing this value by the total sex-averaged autosomal map length of 3146 cM. This can be thought of as a genetic map equivalent to the usual physical map based inbreeding coefficient F_ROH_ (Kardos et al., 2018). This resulted in three inbreeding coefficients per individual, F_ROHlong_, F_ROHmedium_ and F_ROHshort_, each ranging between 0 and 1.

### Genetic simulations

We used simulations to generate baseline expectations for how ROH length classes are expected to differ in their mutation load, and how this is influenced by different selection and dominance coefficients underlying deleterious mutations. Specifically, we used forward genetic Wright-Fisher simulations in SLiM 3 (Haller & Messer, 2019) to simulate deleterious mutations and overlaid neutral mutations using msprime (Kelleher, Etheridge, & McVean, 2016) and pyslim (Haller, Galloway, Kelleher, Messer, & Ralph, 2019).

The Soay sheep were transferred to the St. Kilda archipelago around 4,000 years or roughly 1,000 generations ago (Clutton-Brock & Pemberton, 2004), and their recent Ne has been estimated at 194 (Kijas et al., 2012). We simulated a population with a demographic history close to that estimated for Soay sheep, with a larger ancestral population size N_anc_ = 1,000 for a period of 10,000 generations, followed by an instantaneous change to 200 individuals (after arrival on St. Kilda) for 1,000 generations. Starting 30 generations in the past (at generation 10,970), we simulated an instantaneous bottleneck down to 10 individuals followed by an exponential recovery to 200 individuals within 20 generations to reflect the bottleneck due to the recent transfer of 107 sheep (22 of which were castrates) from the island of Soay to the island of Hirta in 1932, and their rapid population increase to between 600 and 2200 individuals nowadays. This broadly assumes a ratio of effective to census population size of 1:10, which is in line with Soay sheep Ne estimated from genomic data (Kijas et al., 2012)

We modelled 100 Mb diploid genomes with a uniformly distributed recombination rate of 1e-8 per bp per generation, so that the physical distance between two SNPs in Mb was on average equal to their genetic map distance in cM. In each generation, mutations were simulated at a rate of 1e-8 per site, with 30% neutral mutations and 70% deleterious mutations (Kim, Huber, & Lohmueller, 2017). We explored the impact of different parameters underlying the distribution of fitness effects (DFE) for new deleterious mutations by simulating a range of selection and dominance coefficients. Specifically, selection coefficients *s* were drawn from gamma distributions with varying mean *s* ∈ {0.01, −0.03, −0.05} and a shape parameter of 0.2, based on values estimated in humans (Eyre-Walker, Woolfit, & Phelps, 2006). We also varied the dominance coefficients *h* for deleterious alleles from fully to partially recessive with *h ∈* {0, 0.05, 0.2}. SLiM defines a mutation’s fitness effect when homozygous as 1+*s* and when heterozygous as *h*(1+*h*s). Overall, we ran nine simulations for all combinations of *s* and *h,* with 50 replicates each.

At the end of each SLiM simulation, we generated a list of segregating deleterious mutations for the 200 individuals and saved the full tree sequence of the simulation (Haller et al., 2019). Neutral mutations were then added using the coalescent simulator msprime (Kelleher et al., 2016) and pyslim (Haller et al., 2019) and the results for each simulation were saved as vcf files. Before adding neutral mutations, we used recapitation, a technique which runs a coalescent simulation back in time to ensure the coalescence of all samples (Haller et al., 2019). We then called runs of homozygosity in PLINK with the same parameters as in the empirical data analysis above, and clustered ROH into the same three ROH length classes.

Lastly, we combined ROH information with the deleterious mutation data and calculated the following three statistics, all of them as averages across all individuals within a given simulation: 1) The mutation load per cM ROH length. We defined the mutation load per unit length (in cM) for each ROH length class within an individual with 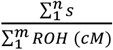, where the numerator sums up the selection coefficients *s* for all *n* deleterious mutations found in the respective ROH class, and the denominator sums up the genetic map lengths of all *m* ROH segments in the relevant class within an individual. This measure of mutation load therefore quantifies the average expected fitness decline per cM of ROH; 2) the average number of deleterious mutations within each ROH class, per cM length; 3) the average allele frequency of deleterious mutations within each ROH class in the population.

### Statistical analyses

To estimate the effects of the three inbreeding coefficients F_ROHlong_,F_ROHmedium_ and F_ROHshort_ on survival, we fitted a binomial Bayesian generalised linear mixed model (GLMM) with logit link, using brms (Bürkner, 2017), a high-level R interface to Stan (Carpenter et al., 2017). The response variable was first-year survival, with a value of 1 if a sheep survived to April 30^th^ in the year after it was born and a value of 0 if it died. We used the following model structure:

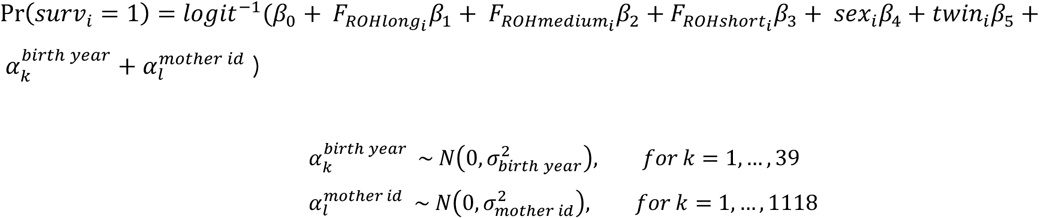

The probability of survival for observation *i* (Pr(*surv_i_* = 1)) was modelled with an intercept *β*_0_, the three population level (fixed) effects for individual inbreeding coefficients F_ROH_ calculated from long ROH (> 12.5 cM), medium ROH (between 1.56 and 12.5 cM) and short ROH (< 1.56 cM) and two further population level effects to take into account the sex of the individual (female = 0, male = 1) and whether it was a twin (no = 0, yes = 1). The model also included two group-level (random) intercept effects for birth year and maternal identity to model environmental variation across years and maternal effects, respectively. The three FROH variables were multiplied by 100, such that the model estimates the change in the odds of survival for a 1% increase in genomic ROH of the respective class. We used a normal prior with mean = 0 and sd = 5 for population-level effects and the default half Student-*t* prior for the standard deviation of group-level parameters. We ran four MCMC chains with the NUTS sampler with 10,000 iterations each, a warmup of 5,000 iterations and no thinning. All chains were visually checked for convergence and the Gelman-Rubin criterion was < 1.1 for all predictors, indicating good convergence (Gelman & Rubin, 1992). We present model estimates as odds ratios, which represent the predicted multiplicative change in the odds of survival for a unit increase in a given predictor, and 95% credible intervals based on the 2.5^th^ and 97.5^th^ percentile in the posterior distribution.

## Results

### ROH in Soay sheep

Overall, we quantified a total of 4,806,614 ROH across all 4,879 Soay sheep, with a mean and maximum genetic map lengths of 1.68 and 80.55 cM, respectively. Individual sheep had on average 625 ROH (range 470-839) spanning 33.3% (range 27.8 – 58.4%) of the autosomal genetic map. Initially, we visually assessed the distribution of ROH lengths over many classes (Supplementary Figure 1), and eventually clustered them into long, medium and short ROH suitable for modelling (Figure 1a). We calculated three individual inbreeding coefficients F_ROHlong_, F_ROHmedium_, and F_ROHshort_ based on these three ROH classes, which varied markedly in their means and distribution in the population (Figure 1 a, Supplementary Figure 1). Long ROH made up only 1.3% of the average Soay sheep genome (mean F_ROHlong_ = 0.013), though the distribution is right skewed and shows that long ROH added up to over 20% of the genome in the most inbred individuals. Medium ROH were the most common class in Soay sheep and made up 21.3% of the average autosomal genome while short ROH made up 10.7% on average.

**Figure 1:**
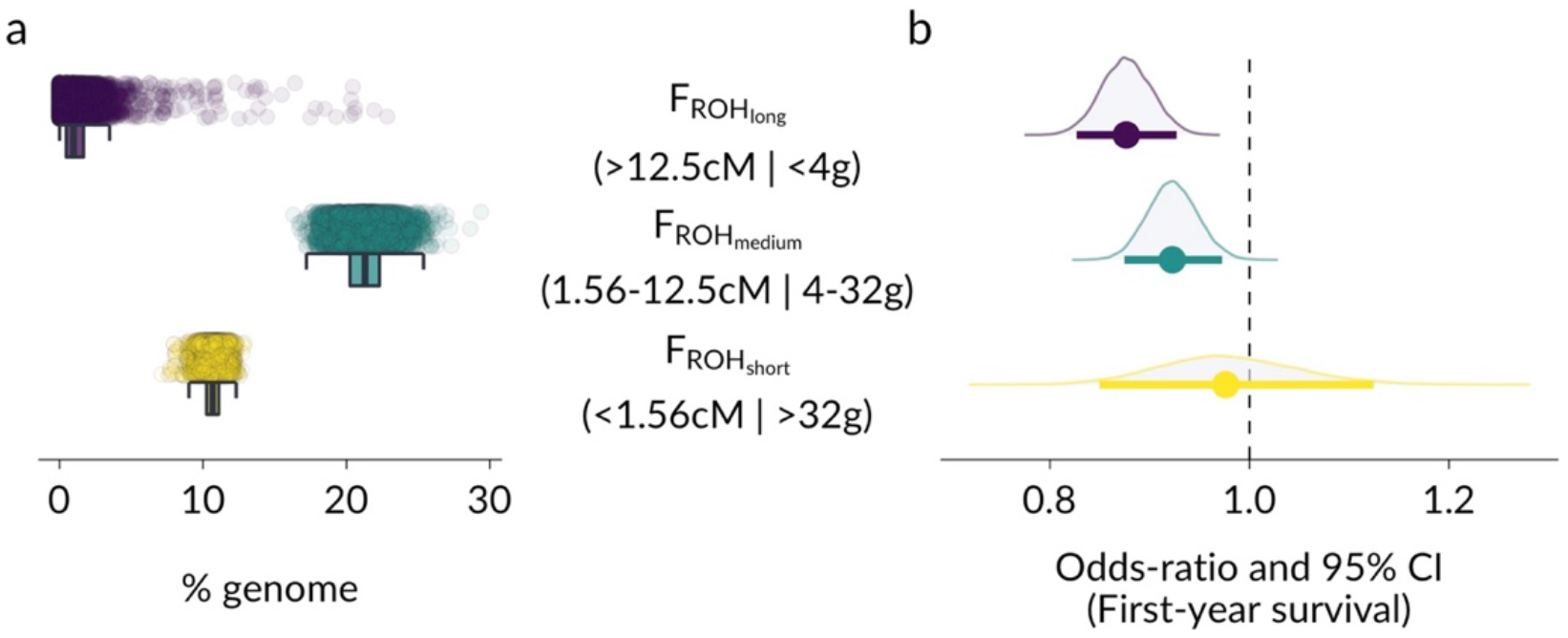
Distribution and fitness effects of inbreeding coefficients F_ROH_ based on different ROH lengths in Soay sheep. Panel a shows the distribution of F_ROHlong_, F_ROHmedium_, and F_ROHshort_ in the population, which were multiplied by 100 to represent the percentage of the genome in the respective ROH length class. Panel b shows the model estimates for the effects of the three inbreeding coefficients on first year survival. The effects are presented as odds-ratios, which show the estimated multiplicative change in the odds of survival for a 1% genomic increase in the respective ROH class. The three classes were clustered by their genetic map length in cM which is associated with the expected time to a MRCA in generations (g).

### Inbreeding depression in survival by ROH length

Inbreeding depression was stronger when F_ROH_ was based on longer ROH (Figure 1b, Supplementary Table 1). The posterior mean odds-ratio (OR) for F_ROHlong_ was 0.876 (95% CI [0.827-0.927]), or an estimated 12.4 % reduction in the odds of survival for a 1% increase in the proportion of the genome found within long ROH. For F_ROHmedium_, the OR was 0.923 (95% CI [0.875-0.973]), corresponding to only a 7.7% reduction in the odds of survival for the same increase in ROH, and F_ROHshort_ were not associated with differences in survival (OR 0.977, 95% CI [0.850-1.125]). In addition, the posterior distributions of the differences in model estimates for F_ROHlong_, F_ROHmedium_, and F_ROHshort_ are also reflecting differences in the estimated effects of inbreeding depression among ROH length classes (Supplementary Figure 2). Lastly, we fitted an alternative model, replacing the three F_ROH_ predictors with the overall inbreeding coefficient F_ROH_ and a second predictor quantifying the mean ROH length per individual. For a given overall inbreeding coefficient F_ROH_, a one cM increase in mean ROH length led to an estimated reduction in the odds of survival by 71% (OR 0.287, 95% CI[0.094, 0.874]), again reflecting stronger inbreeding depression in longer ROH (Supplementary Table 2).

### Genetic simulations

To generate baseline expectations and insights into the reasons for differences in mutation load between ROH length classes we used forward genetic simulations. The overall patterns were qualitatively similar for a range of selection and dominance coefficients for new deleterious mutations (Supplementary Figures 3-5). Therefore, we focus here on the results of simulations based on deleterious mutations following a gamma distribution of fitness effects, with mean *s* = −0.03 and shape parameter *β* = 0.2, where all mutations were partially recessive with a dominance coefficient *h* = 0.05 (Figure 2). Long ROH had the highest overall mutation load per cM length, which was on average 32% lower in medium ROH and 62% lower in short ROH (Figure 2a). The average frequency of deleterious mutations was also lower in long compared with short ROH, showing that longer ROH are enriched for rarer deleterious mutations (Figure 2c). The simulations also reveal a more nuanced pattern: While the overall mutation load per cM was 32% lower in medium compared to long ROH, the average number of deleterious mutations was only 11% lower (Figure 2b). This pattern emerges because rare, strongly deleterious mutations are quickly removed by purifying selection, leading to a substantially lower mutation load in haplotypes originating 4-32 generations ago compared to haplotypes originating less than 4 generations ago. Short ROH (<1.56 cM), with a MRCA more than 32 generations ago had the lowest mutation load and contained substantially fewer deleterious mutation with higher average frequencies (Figure 2).

**Figure 2:**
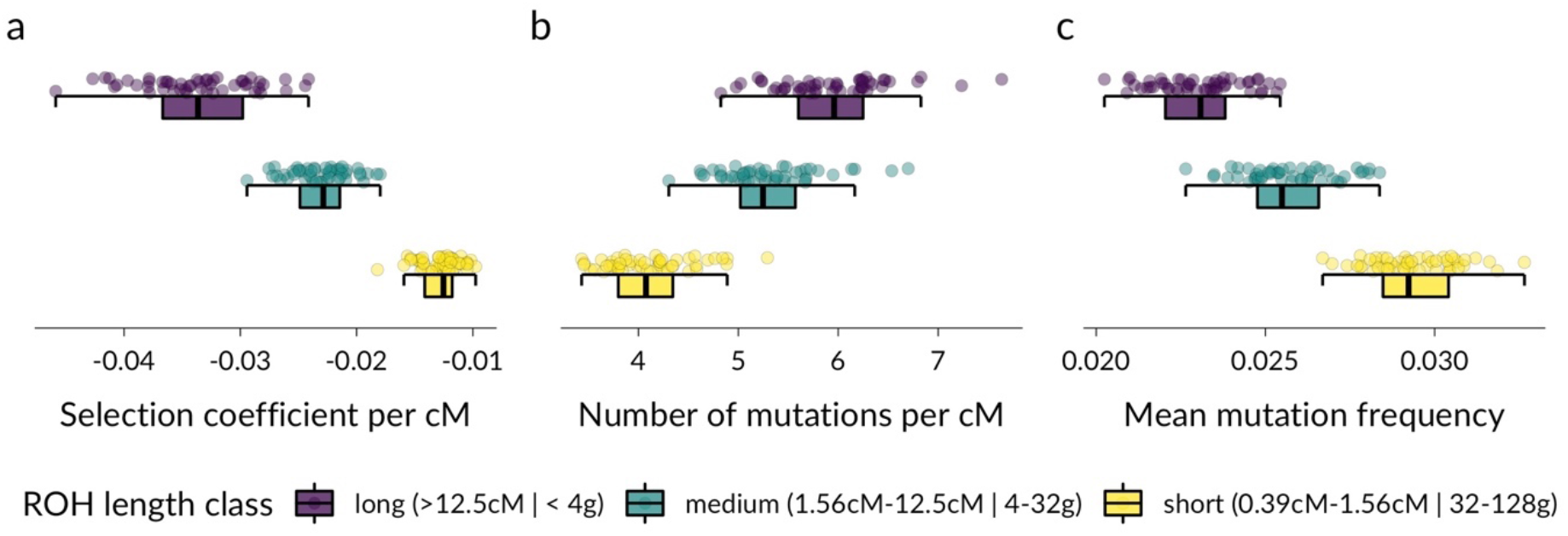
Patterns of deleterious mutations in long, medium and short ROH. Each point represents the mean of 200 individuals of a simulation run. Panel a shows the mean selection coefficient of ROH per cM length, with lower values translating into a larger reduction in individual fitness and therefore representing a higher mutation load. Panel b shows the mean number of deleterious mutations per cM ROH length. Panel c shows the average population frequencies of deleterious mutations.

## Discussion

Long ROH originating from young haplotypes caused stronger inbreeding depression and had a higher mutation load than shorter ROH, which is expected when purifying selection acting over more generations has had more time to purge deleterious variation in older haplotypes. A substantial part of the mutation load is purged within only a few tens of generations, causing a difference in inbreeding depression estimated from medium and long ROH, respectively. Our simulations suggest that this is likely due to selection against strongly deleterious mutations present at low frequencies. While this is theoretically expected in small populations (Hedrick & Garcia-Dorado, 2016; Kimura, Maruyama, & Crow, 1963; Wang, Hill, Charlesworth, & Charlesworth, 1999), empirical evidence based on actual fitness data is rare. However, deleterious mutations can be predicted from genome-sequence data, which has revealed patterns of population-wide purging due to bottlenecks and small population sizes in Mountain Gorillas, Isle Royale Wolves and Alpine Ibex (Grossen et al., 2020; Robinson et al., 2019; Xue et al., 2015). In Soay sheep, the difference in inbreeding depression between long and short ROH despite thousands of years of isolation as a small population suggests that intermediate and strongly deleterious mutations are unlikely to have been completely purged from the population. Instead, these differences probably reflect a haplotype-level snapshot of the ongoing balance between newly arising strongly deleterious mutations expressed in long ROH, and selection against these mutations leaving shorter ROH with lower a mutation load.

Our findings have methodological implications for quantifying inbreeding depression and understanding its genetic architecture. Studies of wild animal population commonly use reduced representation methods such as SNP arrays or RAD sequencing for genotyping individuals. SNP densities might therefore not be high enough to reliably detect short ROH. In Soay sheep, most variation in inbreeding depression was captured by medium and long ROH, which can usually be reliably detected with intermediate SNP densities. In studies of inbreeding depression in wild organisms with low Ne and high linkage disequilibrium, resources might therefore be better allocated into increasing the number of individuals rather than increasing SNP densities from tens of thousands of SNPs to whole-genome-sequencing. When studying the genetic basis of inbreeding depression, ROH can also be used to map the underlying loci in genome-wide association studies (GWAS) (Kardos et al., 2016; Pryce et al., 2014; Stoffel et al., 2020). Our results suggest that the minimum ROH length is important when mapping ROH-fitness relationships. When comparing the fitness of individuals with and without ROH at a given genomic location, the statistical power will be highest when only longer ROH are included, as these are more likely to harbour strongly deleterious recessive alleles. Analyses of the effects of different ROH length classes on fitness prior to GWAS analyses could therefore help to determine an optimal minimum ROH length.

Finally, our analyses provide some fundamental insights into the relationship between deleterious variation, inbreeding depression and purging in a small, wild population. At the haplotype level, we showed that purifying selection constantly removes deleterious variation, causing a difference in the mutation load of IBD haplotypes with different coalescent times. Strongly deleterious mutations are purged relatively quickly, probably because they frequently occur as homozygotes in small populations, which facilitates purging of mutations with large fitness effects despite a relatively low efficiency of selection due to drift (Hedrick & Garcia-Dorado, 2016; Kyriazis, Wayne, & Lohmueller, 2021). Yet, inbreeding depression is strong in Soay sheep, and highly inbred individuals are very unlikely to survive their first winter (Figure 1b, Stoffel et al., 2020). Consequently, inbreeding depression in Soay sheep is probably largely a consequence of the combined effects of many weakly recessive deleterious alleles, which is consistent with a genome-wide association study on the genetic basis of inbreeding depression in Soay sheep (Stoffel et al., 2020). In small populations, theory predicts weakly deleterious mutations will drift more often to higher frequencies and fixation, thereby increasing the mutation load and decreasing mean fitness (Kimura et al., 1963). However, this also decreases the variance in deleterious mutations between individuals and therefore the expected strength of inbreeding depression (Hedrick & Garcia-Dorado, 2016), which is why larger populations are predicted to experience even stronger inbreeding depression than for example Soay sheep. Combining sub-genome level information such as ROH with fitness data is key to assess these theoretical predictions and to gain a deeper understanding of the genetic basis and strength of inbreeding depression in wild populations.

## Acknowledgements

We thank the National Trust for Scotland for permission to work on St. Kilda and QinetiQ, Eurest and Kilda Cruises for logistics and support. We thank Ian Stevenson and many volunteers who have helped with data collection and management and all those who have contributed to keeping the project going. SNP genotyping was conducted at the Wellcome Trust Clinical Research Facility Genetic Core. This work has made extensive use of the Edinburgh Compute and Data Facility (http://www.ecdf.ed.ac.uk/). We are grateful for discussions with Marty Kardos, Jarrod Hadfield, Anna Hewett, Brian Charlesworth and the Wild Evolution Group. We thank Deborah Charlesworth and Emily Humble for detailed comments on earlier versions of the manuscript. The project was funded through an outgoing Postdoc fellowship from the German Science Foundation (DFG) awarded to MAS and a Leverhulme Grant (RPG-2019-072) awarded to JMP and SEJ. Field data collection has been supported by NERC over many years, and most of the SNP genotyping was supported by an ERC Advanced Grant to JMP.

## Author contributions

JMP and MAS designed the study. JGP is the main Soay sheep project fieldworker and collected samples and life history data. JMP has run the Soay sheep long-term study and organised the SNP genotyping. SEJ built the fundamental genomic database, including genotyping, quality control and linkage mapping. MAS conducted data analyses and drafted the manuscript. MAS, JEP and SEJ jointly contributed to concepts, ideas and revisions of the manuscript.

## Data and code accessibility

All data to reproduce the analysis will be uploaded to Dryad upon acceptance. The complete analysis scripts are available on GitHub (https://github.com/mastoffel/sheep_roh).

## References

Aulchenko, Y. S., Ripke, S., Isaacs, A., & Van Duijn, C. M. (2007). GenABEL: An R library for genomewide association analysis. Bioinformatics, 23(10), 1294–1296.

Bérénos, C., Ellis, P. A., Pilkington, J. G., & Pemberton, J. M. (2016). Genomic analysis reveals depression due to both individual and maternal inbreeding in a free-living mammal population. Molecular Ecology, 25(13), 3152–3168.

Browning, S. R., & Browning, B. L. (2015). Accurate non-parametric estimation of recent effective population size from segments of identity by descent. The American Journal of Human Genetics, 97(3), 404–418.

Bürkner, P.-C. (2017). brms: An R package for Bayesian multilevel models using Stan. Journal of Statistical Software, 80(1), 1–28.

Carpenter, B., Gelman, A., Hoffman, M. D., Lee, D., Goodrich, B., Betancourt, M., … Riddell, A. (2017). Stan: A probabilistic programming language. Journal of Statistical Software, 76(1).

Ceballos, F. C., Joshi, P. K., Clark, D. W., Ramsay, M., & Wilson, J. F. (2018). Runs of homozygosity: Windows into population history and trait architecture. Nature Reviews Genetics, 19(4), 220–234. doi: 10.1038/nrg.2017.109

Charlesworth, D., & Willis, J. H. (2009). The genetics of inbreeding depression. Nature Reviews Genetics, 10(11), 783–796. doi: 10.1038/nrg2664

Chen, N., Cosgrove, E. J., Bowman, R., Fitzpatrick, J. W., & Clark, A. G. (2016). Genomic consequences of population decline in the endangered Florida scrub-jay. Current Biology, 26(21), 2974–2979.

Clark, D. W., Okada, Y., Moore, K. H. S., Mason, D., Pirastu, N., Gandin, I., … Wilson, J. F. (2019). Associations of autozygosity with a broad range of human phenotypes. Nature Communications, 10(1), 1–17. doi: 10.1038/s41467-019-12283-6

Clutton-Brock, T. H., & Pemberton, J. M. (2004). Soay sheep: Dynamics and selection in an island population. Cambridge University Press.

Eyre-Walker, A., Woolfit, M., & Phelps, T. (2006). The Distribution of Fitness Effects of New Deleterious Amino Acid Mutations in Humans. Genetics, 173(2), 891–900. doi: 10.1534/genetics.106.057570

Ferenčaković, M., Sölkner, J., Kapš, M., & Curik, I. (2017). Genome-wide mapping and estimation of inbreeding depression of semen quality traits in a cattle population. Journal of Dairy Science, 100(6), 4721–4730. doi: 10.3168/jds.2016-12164

Gelman, A., & Rubin, D. B. (1992). Inference from iterative simulation using multiple sequences. Statistical Science, 7(4), 457–472.

Grossen, C., Guillaume, F., Keller, L. F., & Croll, D. (2020). Purging of highly deleterious mutations through severe bottlenecks in Alpine ibex. Nature Communications, 11(1), 1–12.

Haller, B. C., Galloway, J., Kelleher, J., Messer, P. W., & Ralph, P. L. (2019). Tree-sequence recording in SLiM opens new horizons for forward-time simulation of whole genomes. Molecular Ecology Resources, 19(2), 552–566.

Haller, B. C., & Messer, P. W. (2019). SLiM 3: Forward genetic simulations beyond the Wright-Fisher model. Molecular Biology and Evolution, 36(3), 632–637.

Harrisson, K. A., Magrath, M. J. L., Yen, J. D. L., Pavlova, A., Murray, N., Quin, B., … Sunnucks, P. (2019). Lifetime Fitness Costs of Inbreeding and Being Inbred in a Critically Endangered Bird. Current Biology, 29(16), 2711–2717.e4. doi: 10.1016/j.cub.2019.06.064

Hedrick, P. W., & Garcia-Dorado, A. (2016). Understanding inbreeding depression, purging, and genetic rescue. Trends in Ecology & Evolution, 31(12), 940–952.

Hickey, J. M., Kinghorn, B. P., Tier, B., van der Werf, J. H., & Cleveland, M. A. (2012). A phasing and imputation method for pedigreed populations that results in a single-stage genomic evaluation. Genetics Selection Evolution, 44(1), 9.

Hoffman, J. I., Simpson, F., David, P., Rijks, J. M., Kuiken, T., Thorne, M. A. S., … Dasmahapatra, K. K. (2014). High-throughput sequencing reveals inbreeding depression in a natural population. Proceedings of the National Academy of Sciences of the United States of America, 111(10), 3775–3780. doi: 10.1073/pnas.1318945111

Huisman, J. (2017). Pedigree reconstruction from SNP data: Parentage assignment, sibship clustering and beyond. Molecular Ecology Resources, 17(5), 1009–1024.

Huisman, J., Kruuk, L. E., Ellis, P. A., Clutton-Brock, T., & Pemberton, J. M. (2016). Inbreeding depression across the lifespan in a wild mammal population. Proceedings of the National Academy of Sciences, 113(13), 3585–3590.

Jiang, Y., Xie, M., Chen, W., Talbot, R., Maddox, J. F., Faraut, T., … Zhang, W. (2014). The sheep genome illuminates biology of the rumen and lipid metabolism. Science, 344(6188), 1168–1173.

Johnston, S. E., Bérénos, C., Slate, J., & Pemberton, J. M. (2016). Conserved Genetic Architecture Underlying Individual Recombination Rate Variation in a Wild Population of Soay Sheep (Ovis aries). Genetics, 203(1), 583–598. doi: 10.1534/genetics.115.185553

Kardos, M., Åkesson, M., Fountain, T., Flagstad, Ø., Liberg, O., Olason, P., … Ellegren, H. (2018). Genomic consequences of intensive inbreeding in an isolated wolf population. Nature Ecology & Evolution, 2(1), 124–131. doi: 10.1038/s41559-017-0375-4

Kardos, M., Luikart, G., & Allendorf, F. W. (2015). Measuring individual inbreeding in the age of genomics: Marker-based measures are better than pedigrees. Heredity, 115(1), 63–72. doi: 10.1038/hdy.2015.17

Kardos, M., Qvarnström, A., & Ellegren, H. (2017). Inferring individual inbreeding and demographic history from segments of identity by descent in Ficedula flycatcher genome sequences. Genetics, 205(3), 1319–1334.

Kardos, M., Taylor, H. R., Ellegren, H., Luikart, G., & Allendorf, F. W. (2016). Genomics advances the study of inbreeding depression in the wild. Evolutionary Applications, 9(10), 1205–1218.

Kelleher, J., Etheridge, A. M., & McVean, G. (2016). Efficient coalescent simulation and genealogical analysis for large sample sizes. PLoS Computational Biology, 12(5), e1004842.

Kijas, J. W., Lenstra, J. A., Hayes, B., Boitard, S., Neto, L. R. P., Cristobal, M. S., … Consortium, other members of the I. S. G. (2012). Genome-Wide Analysis of the World’s Sheep Breeds Reveals High Levels of Historic Mixture and Strong Recent Selection. PLOS Biology, 10(2), e1001258. doi: 10.1371/journal.pbio.1001258

Kim, B. Y., Huber, C. D., & Lohmueller, K. E. (2017). Inference of the Distribution of Selection Coefficients for New Nonsynonymous Mutations Using Large Samples. Genetics, 206(1), 345–361. doi: 10.1534/genetics.116.197145

Kimura, M., Maruyama, T., & Crow, J. F. (1963). The Mutation Load in Small Populations. Genetics, 48(10), 1303–1312.

Kyriazis, C. C., Wayne, R. K., & Lohmueller, K. E. (2020). Strongly deleterious mutations are a primary determinant of extinction risk due to inbreeding depression. BioRxiv, 678524. doi: 10.1101/678524

Kyriazis, C. C., Wayne, R. K., & Lohmueller, K. E. (2021). Strongly deleterious mutations are a primary determinant of extinction risk due to inbreeding depression. Evolution Letters, 5(1), 33–47. doi: https://doi.org/10.1002/evl3.209

Morrissey, M. B., Parker, D. J., Korsten, P., Pemberton, J. M., Kruuk, L. E., & Wilson, A. J. (2012). The prediction of adaptive evolution: Empirical application of the secondary theorem of selection and comparison to the breeder’s equation. Evolution: International Journal of Organic Evolution, 66(8), 2399–2410.

Niskanen, A. K., Billing, A. M., Holand, H., Hagen, I. J., Araya-Ajoy, Y. G., Husby, A., … Jensen, H. (2020). Consistent scaling of inbreeding depression in space and time in a house sparrow metapopulation. Proceedings of the National Academy of Sciences, 117(25), 14584–14592. doi: 10.1073/pnas.1909599117

Pemberton, T. J., & Szpiech, Z. A. (2018). Relationship between Deleterious Variation, Genomic Autozygosity, and Disease Risk: Insights from The 1000 Genomes Project. The American Journal of Human Genetics, 102(4), 658–675. doi: 10.1016/j.ajhg.2018.02.013

Pryce, J. E., Haile-Mariam, M., Goddard, M. E., & Hayes, B. J. (2014). Identification of genomic regions associated with inbreeding depression in Holstein and Jersey dairy cattle. Genetics Selection Evolution, 46(1). doi: 10.1186/s12711-014-0071-7

Purcell, S., Neale, B., Todd-Brown, K., Thomas, L., Ferreira, M. A., Bender, D., … Daly, M. J. (2007). PLINK: A tool set for whole-genome association and population-based linkage analyses. The American Journal of Human Genetics, 81(3), 559–575.

Ralls, K., Sunnucks, P., Lacy, R. C., & Frankham, R. (2020). Genetic rescue: A critique of the evidence supports maximizing genetic diversity rather than minimizing the introduction of putatively harmful genetic variation. Biological Conservation, 251, 108784. doi: 10.1016/j.biocon.2020.108784

Robinson, J. A., Brown, C., Kim, B. Y., Lohmueller, K. E., & Wayne, R. K. (2018). Purging of strongly deleterious mutations explains long-term persistence and absence of inbreeding depression in island foxes. Current Biology, 28(21), 3487–3494.

Robinson, J. A., Räikkönen, J., Vucetich, L. M., Vucetich, J. A., Peterson, R. O., Lohmueller, K. E., & Wayne, R. K. (2019). Genomic signatures of extensive inbreeding in Isle Royale wolves, a population on the threshold of extinction. Science Advances, 5(5), eaau0757.

Stoffel, M. A., Johnston, S. E., Pilkington, J. G., & Pemberton, J. M. (2020). Genetic architecture and lifetime dynamics of inbreeding depression in a wild mammal. BioRxiv.

Szpiech, Z. A., Mak, A. C., White, M. J., Hu, D., Eng, C., Burchard, E. G., & Hernandez, R. D. (2018). Ancestry-dependent enrichment of deleterious homozygotes in runs of homozygosity. BioRxiv, 382721.

Szpiech, Z. A., Xu, J., Pemberton, T. J., Peng, W., Zöllner, S., Rosenberg, N. A., & Li, J. Z. (2013).Long runs of homozygosity are enriched for deleterious variation. The American Journal of Human Genetics, 93(1), 90–102.

Thompson, E. A. (2013). Identity by descent: Variation in meiosis, across genomes, and in populations. Genetics, 194(2), 301–326.

Wang, J., Hill, W. G., Charlesworth, D., & Charlesworth, B. (1999). Dynamics of inbreeding depression due to deleterious mutations in small populations: Mutation parameters and inbreeding rate. Genetics Research, 74(2), 165–178. doi: 10.1017/S0016672399003900

Xue, Y., Prado-Martinez, J., Sudmant, P. H., Narasimhan, V., Ayub, Q., Szpak, M., … Cooper, D. N. (2015). Mountain gorilla genomes reveal the impact of long-term population decline and inbreeding. Science, 348(6231), 242–245.

