## Supplementary_Figures_and_Tables for "Mutation load decreases with haplotype age in wild Soay sheep"

Authors names and addresses:

Stoffel, M.A.[1*], Johnston, S.E.[1], Pilkington, J.G.[1], Pemberton, J.M[1]

[1]Institute of Evolutionary Biology, School of Biological Sciences, University of Edinburgh, Edinburgh, EH9 3FL, United Kingdom

* Corresponding author:
Martin A. Stoffel
Postal address: Institute of Evolutionary Biology, University of Edinburgh, Edinburgh, EH9 3FL, UK

### Supplementary Figures

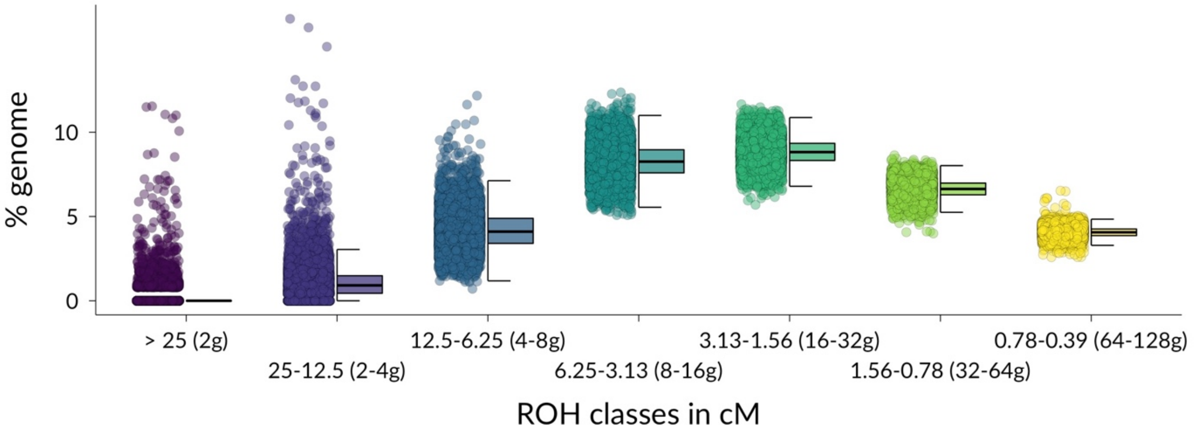


**Supplementary Figure 1: Distribution of different ROH lengths classes in Soays heep**. ROH were measured in cM and clustered by their expected time to most recent common ancestor ranging from 2 to 128 generations ago. Each point represents the proportion of ROH of a specific length class in the genome of an individual.

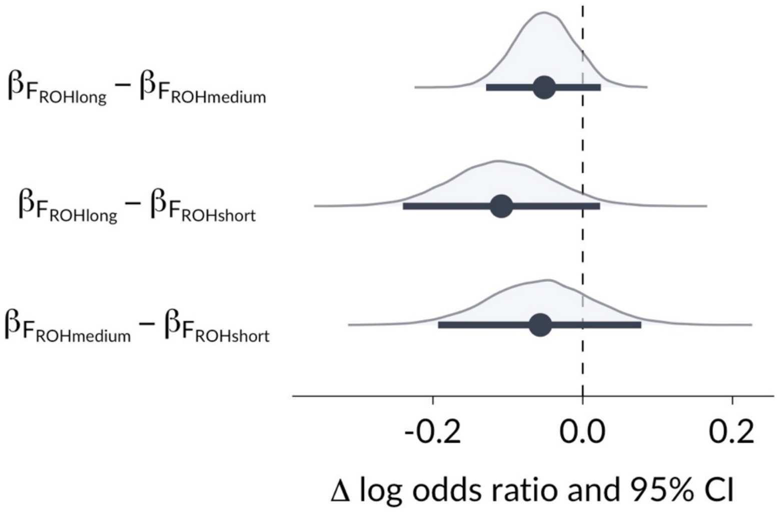


**Supplementary Figure 2: Posterior distribution for the differences in effect size estimates for inbreeding depression due to long, medium and short ROH**. In contrast to Figure 2a, the differences in model estimates are shown on the untransformed log-odds (logit) scale, which facilitates visualising the differences in model estimates for inbreeding depression based on the three different inbreeding coefficients. The posterior distributions are largely negative, indicating inbreeding depression estimates based longer ROH tended to have larger (negative) effect sizes.

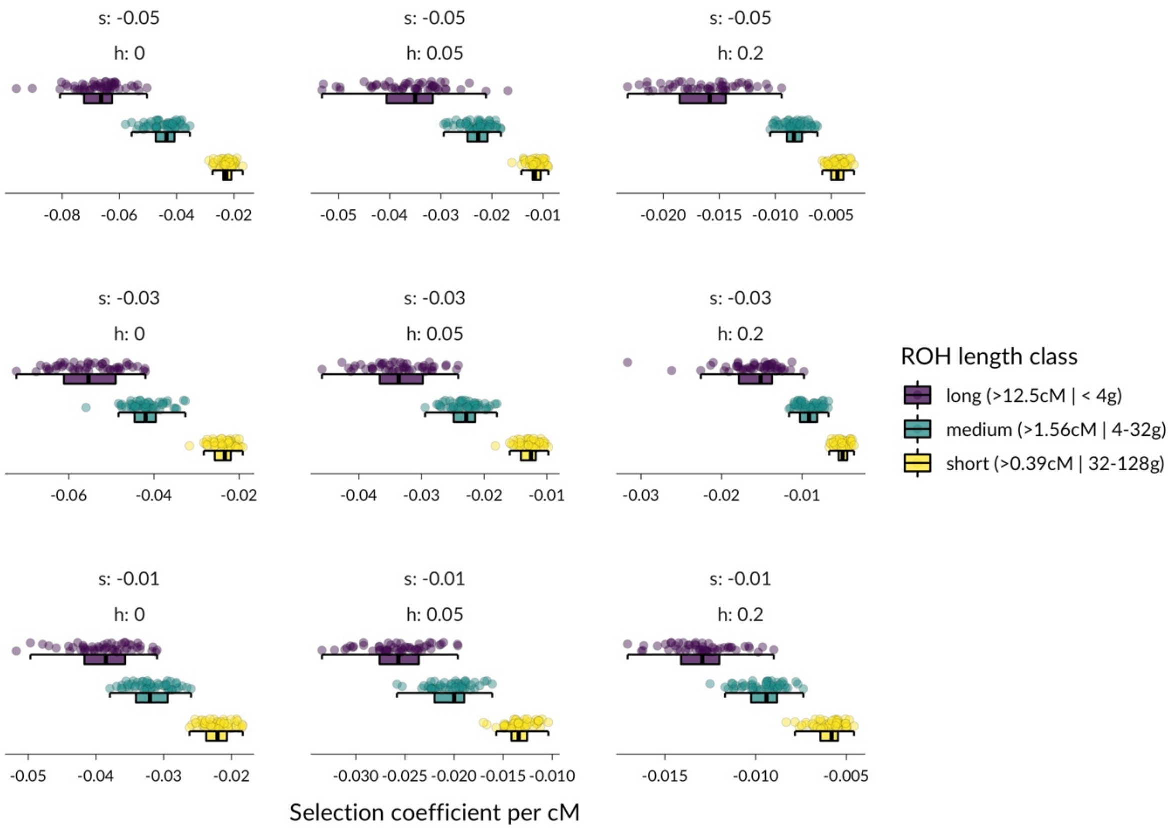


**Supplementary Figure 3: Mutation load (selection coefficient per cM) within ROH length classes for different DFE parameters**. Each datapoint represents the average selection coefficient per cM for three ROH length classes, calculated as the mean across 200 individuals sampled at the end of each simulation. The selection coefficients for new deleterious mutations were drawn from gamma distributions with mean s ∈ {-0.01, -0.03, -0.05} and shape 0.2 and with three different dominance coefficients h ∈ {0, 0.05, 0.2}. The results of simulations with each combination of s and h are shown in the nine panels.

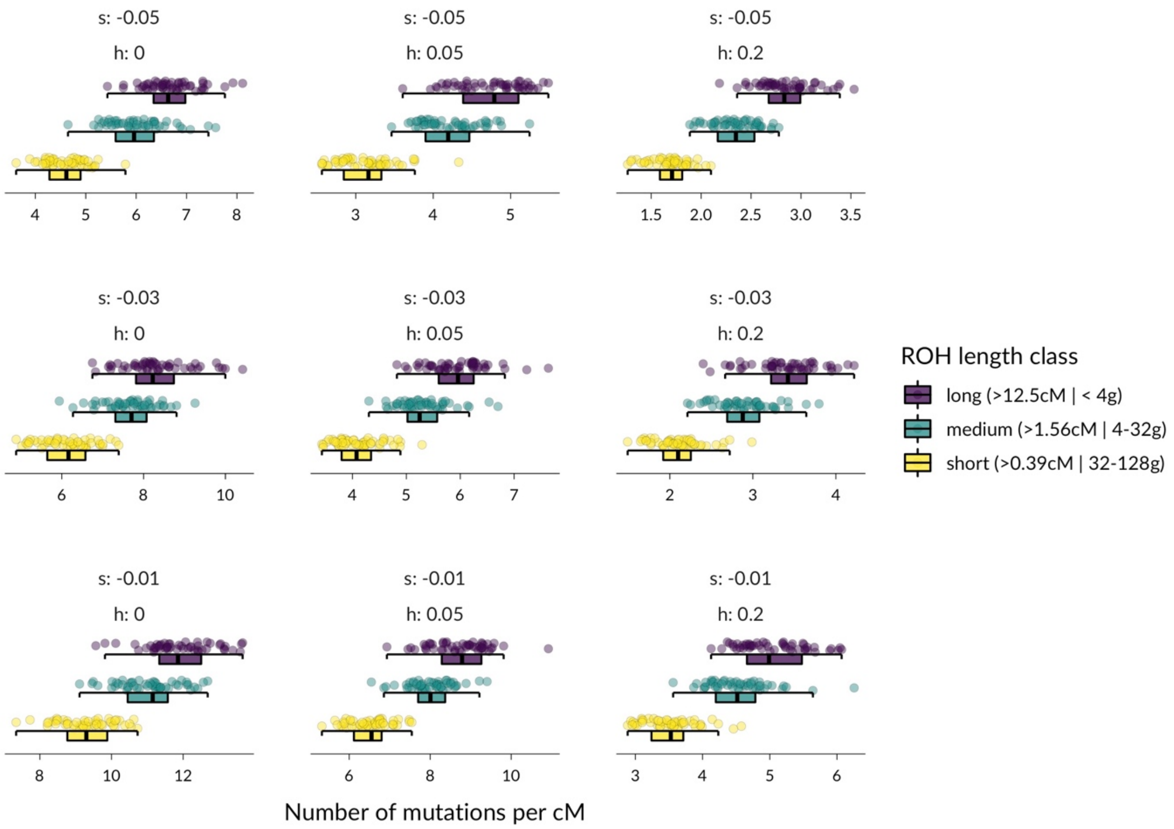


**Supplementary Figure 4: Abundance of deleterious mutations within ROH length classes for different DFE parameters**. Each datapoint represents the number of deleterious mutations per cM for three ROH length classes, calculated as the mean across 200 individuals sampled at the end of each simulation. The selection coefficients for new deleterious mutations were drawn from gamma distributions with mean s ∈ {-0.01, -0.03, -0.05} and shape 0.2 and with three different dominance coefficients h ∈ {0, 0.05, 0.2}. The results of simulations with each combination of mean s and h are shown in the nine panels.

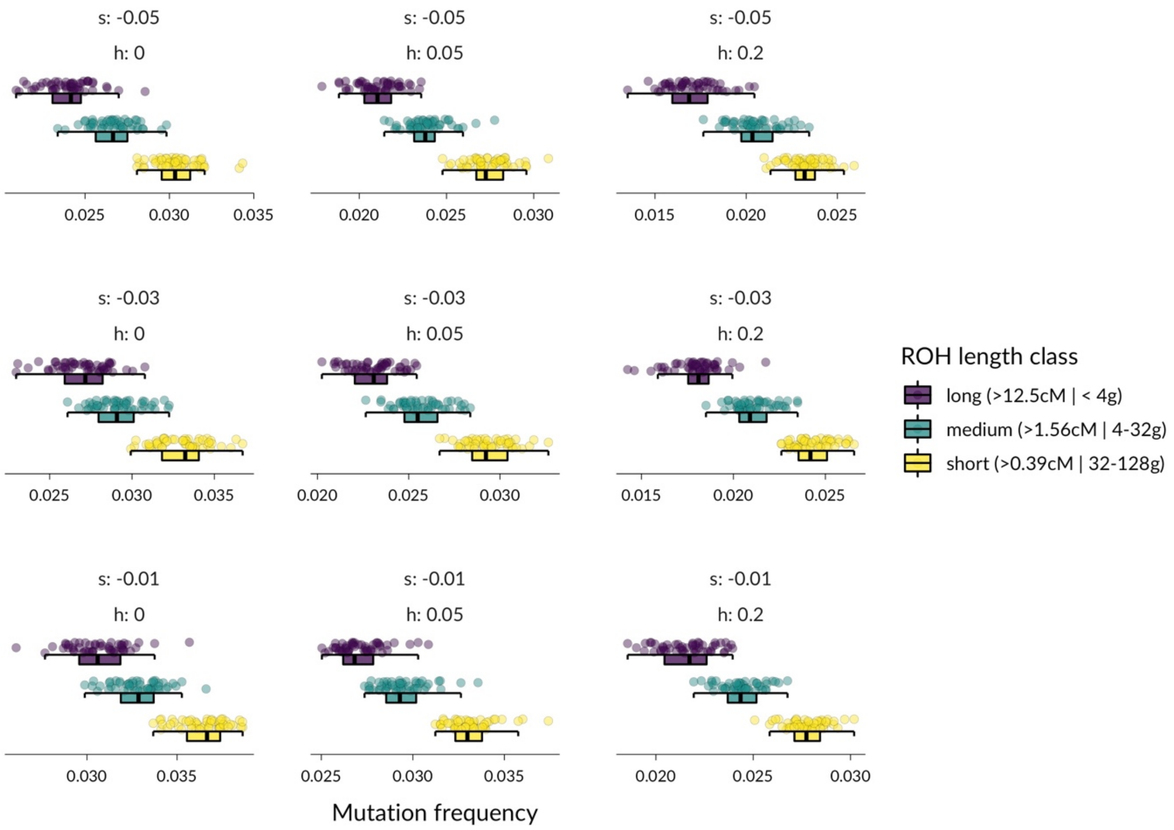


**Supplementary Figure 5: Deleterious mutation frequencies within ROH length classes for different DFE parameters**. Each datapoint represents the mean frequency of deleterious mutations contained in three ROH length classes, calculated as the mean across 200 individuals sampled at the end of each simulation. The selection coefficients for new deleterious mutations were drawn from gamma distributions with mean s ∈ {-0.01, -0.03, -0.05} and shape 0.2 and with three different dominance coefficients h ∈ {0, 0.05, 0.2}. The results of simulations with each combination of mean s and h are shown in the nine panels.

#### Supplementary Tables

| Term | Post.Mean | CI (2.5%) | CI (97.5%) | Info |
|---|---|---|---|---|
| Intercept | 2.423 (11.281) | 0.214 (1.239) | 4.658 (105.385) | |
| Population level/fixed effects | | | | |
| $F_{ROHlong}$ | -0.132 (0.876) | -0.19 (0.827) | -0.076 (0.927) | continuous |
| $F_{ROHmedium}$ | -0.081 (0.923) | -0.134 (0.875) | -0.028 (0.973) | continuous |
| $F_{ROHshort}$ | -0.024 (0.977) | -0.163 (0.85) | 0.117 (1.125) | continuous |
| Sex | -0.67 (0.512) | -0.825 (0.438) | -0.519 (0.595) | categorical (0=male, 1=female) |
| Twin | -1.027 (0.358) | -1.242 (0.289) | -0.811 (0.445) | categorical (0=singleton, 1=twin) |
| Group level/random effects (standard deviation) | | | | |
| Birth year | 1.992 (7.33) | 1.501 (4.486) | 2.661 (14.306) | n = 1118 |
| Mother ID | 0.723 (2.061) | 0.585 (1.795) | 0.863 (2.369) | n = 39 |

**Supplementary Table 1: Model estimates for a Bayesian animal model of annual survival with binomial error structure and logit link.** Shown are the model estimates for the posterior mean and the lower and upper credible interval using the 2.5$^{th}$ and 97.5$^{th}$ quantile on both the logit scale and the odds-ratio scale in round brackets. The three $F_{ROH}$ predictors have been multiplied by 100, so that the model directly estimates the log-odds change in survival for a 1% increase in genomic ROH. The last column shows the reference levels for the categorical predictors and the number of groups for group-level effects.

| Term | Post.Mean | CI (2.5%) | CI (97.5%) | Info |
|---|---|---|---|---|
| Intercept | 3.953 (52.108) | 2.521 (12.437) | 5.409 (223.51) | |
| Population level/fixed effects | | | | |
| $F_{ROH}$ | -0.047 (0.954) | -0.114 (0.892) | 0.02 (1.02) | continuous |
| Mean ROH length (cM) | -1.248 (0.287) | -2.365 (0.094) | -0.135 (0.874) | continuous |
| Sex | -0.67 (0.512) | -0.823 (0.439) | -0.516 (0.597) | categorical (0=male, 1=female) |
| Twin | -1.026 (0.359) | -1.242 (0.289) | -0.813 (0.444) | categorical (0=singleton, 1=twin) |
| Group level/random effects (standard deviation) | | | | |
| Birth year | 1.988 (7.3) | 1.496 (4.463) | 2.655 (14.221) | n = 1118 |
| Mother ID | 0.717 (2.048) | 0.578 (1.782) | 0.855 (2.352) | n = 39 |

**Supplementary Table 2: Model estimates for an alternative Bayesian animal model of annual survival with binomial error structure and logit link.** Shown are the model estimates for the posterior mean and the lower and upper credible interval using the 2.5$^{th}$ and 97.5$^{th}$ quantile on both the logit scale and the odds-ratio scale in round brackets. The three $F_{ROH}$ predictors have replaced by an overall inbreeding coefficient $F_{ROH}$ including all ROH and a predictor quantifying the mean length of ROH within an individual. The negative fitness of ROH length shows that when keeping $F_{ROH}$ constant, individuals with longer ROH have a lower survival probability. Note that $F_{ROH}$ and Mean ROH length correlate highly (r = 0.85), which is why the estimate for $F_{ROH}$ is relatively low. In addition, $F_{ROH}$ has been multiplied by 100 again, so that the log-odds ratio and odds-ratio estimate a change in survival probability for a 1% increase in ROH.
